# Hierarchical Network Exploration using Gaussian Mixture Models

**DOI:** 10.1101/623157

**Authors:** James Mathews, Saad Nadeem, Maryam Pouryahya, Zehor Belkhatir, Joseph O. Deasy, Allen Tannenbaum

## Abstract

We present a framework based on optimal mass transport to construct, for a given network, a reduction hierarchy which can be used for interactive data exploration and community detection. Given a network and a set of numerical data samples for each node, we calculate a new computationally-efficient comparison metric between Gaussian Mixture Models, the Gaussian Mixture Transport distance, to determine a series of merge simplifications of the network. If only a network is given, numerical samples are synthesized from the network topology. The method has its basis in the local connection structure of the network, as well as the joint distribution of the data associated with neighboring nodes.

The analysis is benchmarked on networks with known community structures. We also analyze gene regulatory networks, including the PANTHER curated database and networks inferred from the GTEx lung and breast tissue RNA profiles. Gene Ontology annotations from the EBI GOA database are ranked and superimposed to explain the salient gene modules. We find that several gene modules related to highly specific biological processes are well-coordinated in such tissues. We also find that 18 of the 50 genes of the PAM50 breast-tumor prognostic signature appear among the highly coordinated genes in a single gene module, in both the breast and lung samples. Moreover these 18 are precisely the subset of the PAM50 recently identified as the basal-like markers.

## 1 Introduction

The growing importance of complex networks has been documented in a huge and growing literature, now being referred to as the field of *network science* [2]. A key problem is the representation of network data in a readily accessible format. Ideally the representation should be amenable to human-in-the-loop, interactive, exploratory data analysis.

Supernode methods have been used previously to compress very large networks to a desired level of resolution, mainly toward the goal of improving the computational performance of community detection algorithms [10, 14]. The key idea of supernodes is to group nodes into modules and consider the new network comprised of the connections between groups implied by the individual node connections. In its simplest form, the groups may be obtained by devising an edge weighting intended to measure similarity between neighboring nodes, and successively “collapsing” edges beginning with the greatest similarity to create a hierarchical representation.

A natural improvement is to find approximations of a given network “from below” with gradually expanding small node subsets, an approach developed by Stanley *et al*. [10]. The authors randomly distribute seed nodes which are then expanded into supernodes using direct neighborhoods. In a more global approach, Yang *et al*. [14] present a method of supernode network representation involving explicit consideration of known prior constraints on the set of network topologies of interest for a low-complexity approximation to the given network satisfying the constraints. For a more detailed survey on this topic, refer to Besta *et al*. [3].

Unlike the method of [10] and in common with the method of [14], our proposed approach makes essential use of additional data beyond the network topology. Rather than qualitative constraints as in [14], we assume numerical data along the nodes of the network. Data of this type is frequently available in practice when each node represents a variable or feature of interest across a sample set. For example, RNA expression of a gene (node) across a patient tissue sample population. Such data can also be generated or synthesized from the underlying network topology if the topology is the primary variable of interest and sample analysis is not needed. Conversely the network topology may be inferred from the sample data if no prior network is known. The node weightings are interpreted as defining samples from the joint distribution of random variables associated with the nodes.

A very natural model for distributions is the Gaussian mixture, used in many data processing and analysis applications [8]. In general, a ***mixture model*** is a weighted linear combination of distributions where each component represents a subpopulation. In particular, the Gaussian mixture model (GMM) is a weighted average of Gaussians. GMMs are popular given their versatility and overall simplicity in data representation. They are ubiquitous in statistics, hypothesis testing, decision theory, and machine learning. The idea is that real-world data may not be densely distributed on a high dimensional space, and instead is concentrated in a low dimensional subspace. Further, in many cases of interest, the data is sparsely distributed into a number of subgroups, and so differences within a given subgroup are not as important as those among the subgroups. Mixture models capture these properties, and this motivated the work of Chen *et al*. [4] modifying optimal mass transport (OMT) theory [12, 13] into a form suitable for Gaussian mixture models.

OMT provides the Gaussian mixture framework with a natural comparison metric between mixtures, and reciprocally mixtures provide a natural model with which to make the computation of OMT tractable. We use GMMs to model the functional role played by a node with respect to the data along its neighbors in the network. This role is quantified by the average GMM/OMT distance between two nodes, which we call ***Gaussian Mixture Transport*** (GMT) distance. Hierarchical clustering then provides a simplified version of the network for each given level of complexity. The simplified or compressed network represents a “projection” of the prior network which is most relevant according to the evidence observed in the data.

## 2 Methods

To illustrate the construction of GMT distance-based network reduction hierarchies, we show the steps of the construction applied to a concrete example, a network synthesized in order to have a known community structure. The edge density is high within the communities, and low between communities. Node weightings are randomly generated in terms of the network topology by neighbor averaging or the iterated graph Laplacian of random weightings, with one example of such a weighting depicted in Figure 2 (c). A preview of the series of network simplifications is shown in Figure 1.

**Figure 1:**
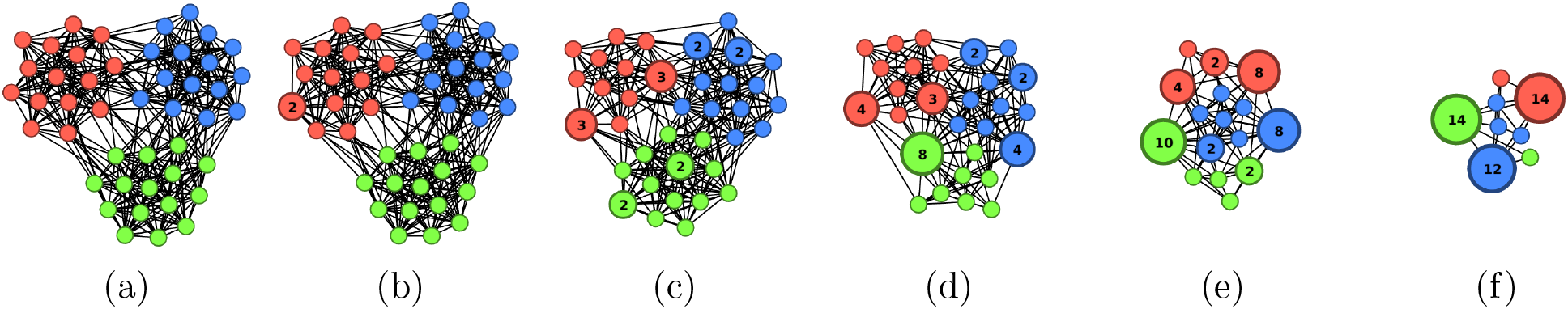
A synthetic network with K=3 communities containing N=45 nodes total. The network was randomly generated to have within-community edge connectivity 0.08 out of a possible maximum of 0.68, and between-community edge connectivity 0.28 out of a possible maximum of 0.32. A random node weighting can be generated from the network by iterated neighborhood-averaging applied to an initial node weighting equal to 1 on a randomly selected node and 0 on every other node. 200 node weightings were generated in this manner. Hierarchical clustering was performed with respect to GMT-distance similarity edge scores. (a) The original network. (b)-(f) 5 selected hierarhical levels from the series. Each node group is labelled by the number of nodes it contains.

**Figure 2:**
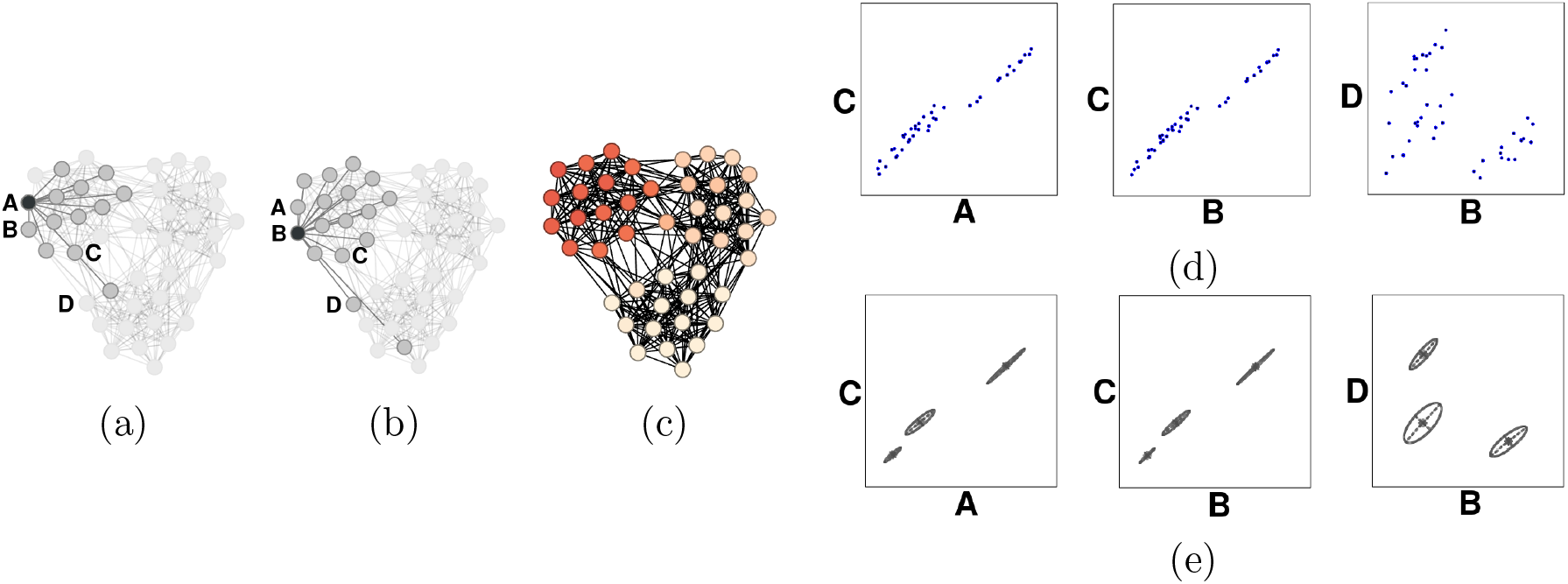
Illustration of the calculation of the GMT distance for selected nodes in the network depicted in Figure 1. Nodes labelled A and B were selected from the same community. (a) The neighborhood of node A. (b) The neighborhood of node B. Both of these neighborhoods contain node C, while node D is only in the neighborhood of B. (c) One of the 200 randomly generated node weightings described in Figure 1. (d) The joint distributions of the weight values for node pairs (C,A), (C,B), and (D,B). (e) The 3-population 2D Gaussian Mixture Models for these joint distributions. The GMM/OMT distance between model (C,A) and model (C,B) is 3.3 x 10^−7^. This value is typical among models (C′, A) and (C′, B) where C′ is any other node besides C intervening between A and B, so the GMT distance between A and B, which averages these values, will be small. This reflects the fact that A and B belong to a highly connected community. By contrast the GMM/OMT distance between models (C,B) and (C,D) is 1.0 x 10^−4^, a value typical among C′ intervening between B and D. So the GMT distance between B and D will be relatively large, reflecting the fact that B and D were selected from different highly connected communities. In this example it is not essential that C was selected from the same community as A and B; though the typical mutual neighbor of A and B will be contained in the same community, like C, the same conclusions follow when C is selected from elsewhere in the network.

### 2.1 Gaussian Mixture Transport distance

A naive approach would group nodes together based on similar properties, for example, by making comparisons between the (univariate) distributions of the weight data associated with each node. Comparison in this case is a classical topic, addressed, for instance, by the Kolmogorov-Smirnov test. One step beyond this is to summarize the joint (bivariate) distribution associated with each edge node-pair by a Pearson correlation or related metric, and then use this similarity metric for classical clustering.

For a higher-order approach, we consider the bivariate distributions associated with each edge (the joint distributions of the variables associated with the two endpoint nodes), then compare *pairs* of bivariate distributions associated with adjacent edges A-C and B-C (in this case, adjacent along node C). If the distributions are similar, A and B will be considered to have a similar “functional role” in the network locally near C. Ultimately we will summarize this similarity over all C intermediate between A and B.

How is the similarity between bivariate distributions quantified? The Bhattacharyya distance [5] is one similarity measure between distributions in higher dimensions. However, a direct calculation of this distance tends to require rasterization of the space involved and may be prohibitively costly to compute. The theory developed by Chen *et al*. [4] for Gaussian Mixture Models provides a nearly closed-form alternative which is well-suited to the task, a metric we call Gaussian Mixture Transport similarity. First, the distributions are approximated by Gaussian mixtures with a given number of subpopulations. The mixture weights are interpreted as probability distributions on the discrete set of subpopulations, which are themselves compared using the optimal mass transport metric or Earth Mover’s Distance (EMD). For this calculation of the EMD, the cost function corresponding to motion from the discrete point labelling a subpopulation of the first mixture to a discrete point labelling a subpopulation of the second mixture is taken to be the actual optimal mass transport distance between the corresponding ordinary Gaussian distributions. The GMT distance between two nodes (not necessarily connected by an edge) is this GMM/OMT distance averaged over all adjacent edges with the same free endpoints. For a detailed description, see Algorithm 2 summarizing the formulae of [4] and the OMT Background section A. An example of the GMT distance comparison of node weight distributions is illustrated in Figure 2. The last step is classical hierarchical clustering using the GMT distance sparse similarity matrix. We use the average-distance-based hierarchical clustering method, though the standard alternatives single-linkage, complete-linkage, Ward clustering, etc. may be used depending on the application.

### 2.2 Implementation and runtime complexity

A naive version of Algorithm 1 would iterate over all edge pairs, with complexity class O(*E*^2^) where *E* is the number of edges of the network. However, since only adjacent edge pairs are used, we instead iterate over the nodes and then over the pairs of its neighbors, with complexity class O(*N D*^2^) where *N* is the number of nodes and *D* is the maximum degree over all nodes.

The mixture modeling and GMT distance calculations are classically parallelizable. The mixture modeling because it only depends on the initial node variable pair distributions, and the GMT distances because they only depend on the resulting list of mixture models. This makes our algorithm feasible for rapid computation. The discrete Earth Mover’s Distance is performed with the R package ‘emdist’ [11]. The mixture modeling itself is performed with the R package ‘mclust’ [9]. In practice, the number *P* of mixture model populations has little effect on the overall output and performance as long as *P* lies in the approximate range from 3 to 10. If the number of node weighting samples is as low as a few hundred, it is not meaningful to choose *P* much greater than 10 anyway, since the number of data points per “population” should not be too low. High accuracy of the mixture model as a representation of the joint distribution of two given node variables is not essential for the purpose of inferring distances between the distributions from distances between the models.

### 2.3 Visualization

Once the hierarchy is computed, it is formatted for viewing in the Gephi graph visualization software using a custom plugin. One view shows the original feature network with some new edges with weights chosen to highlight a given hierarchical level; the weights are calculated as a sigmoid function of GMT distance matrix values. The level can be selected in real-time.

For larger graphs, a static view is used to reduce the computational burden of real-time rendering. For this we use the hierarchy itself considered as a graph. We note that this visualization method applies to any hierarchical clustering and could serve as a general-purpose alternative to the usual rectilinear branch representation often used to decorate heatmaps. The graph is a tree or union of trees, with leaf nodes representing features and internal nodes representing feature groups. We choose leaf node sizes to reflect the linkage height, or hierarchical level, of the first internal node to which it is attached. Lower levels correspond to larger nodes, since nodes joining the tree at a lower level do so on the basis of stronger evidence of coordination with other nodes (lower GMT distances), which we wish to highlight. Internal nodes are given negligible size. A planar representation with no edge-crossings is possible since the graph consists of trees. A force-directed layout is used to arrange the nodes in a way guided by the tree structure and node sizes.

## 3 Benchmarks

Although the GMT hierarchy is primarily designed for reducing the feature structure of a numerical dataset, it can also be applied to a pure network topology by means of synthesized node weights. For this we use the iterated graph Laplacian Δ applied to single-node weightings with randomly chosen support nodes. The resulting weightings can be understood as random linear combinations of the Δ eigenfunctions, with emphasis on those eigenfunctions with large eigenvalues. The spectrum of Δ is well-studied and known to capture a lot of detailed information about the underlying graph. See Figure 4 for example scatter plots of this type of data, synthesized from the PANTHER curated gene network.

The hierarchy can be completed to an unsupervised community detection algorithm using a numerical metric of modularity, in the usual way, by selecting from among the level-cutoff clusterings the one with the best value of the metric. In the supervised setting, the level cutoff can be selected to give the best value of a cluster similarity metric like Normalized Mutual Information (NMI) calculated against a known community structure. Figure 3 compares the GMT community detection method with three established methods available in the R ‘igraph’ library: greedy optimization, Louvain optimization, and label propagation.

**Figure 3:**
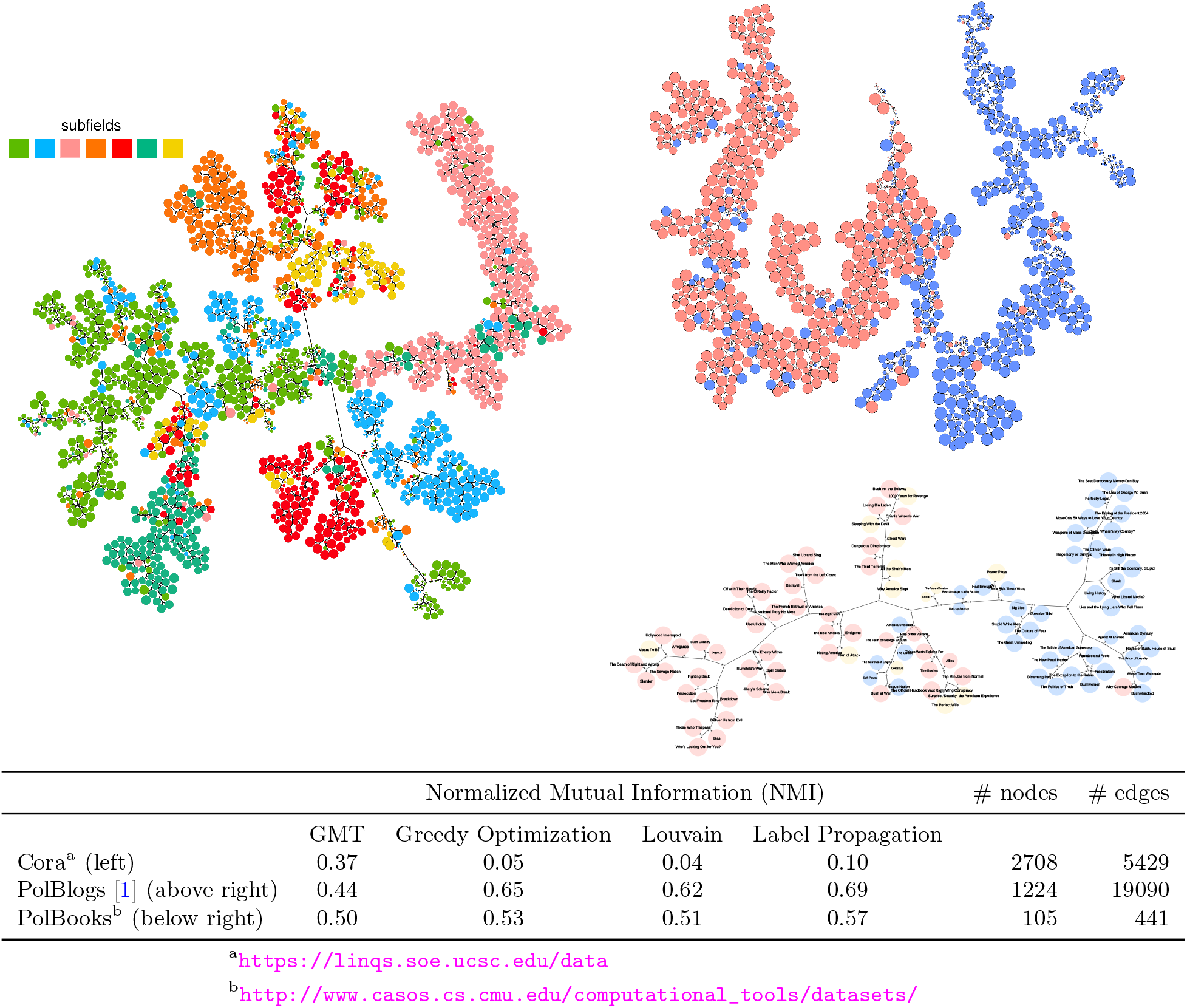
Comparison of GMT community detection with greedy optimization of the modularity, Louvain optimization, and label propagation, with respect to NMI. PolBlogs and PolBooks are respectively a network of political blog links and a copurchasing network for political books in the United States. Both are provided with manually annotated indications of political leaning for each node (red and blue). For these smaller networks with few, well-defined communities, the established methods outperform GMT. On the larger Cora academic paper citation network, presumably with greater real-world complexity, GMT outperforms the other methods by a factor of 3.7. The visualization substantially reveals the manually annotated subfields, and also seems to suggest an improvement where some putative subfields are divided into multiple distinct groups and closely related to specific alternative subfields. For example, the subfield in red is divided into two tightly clustered groups, one group very close to the subfields in orange and yellow.

## 4 Analysis of Gene Regulatory Networks

Gene and protein networks are now ubiquitous in medicine and biology. Though they are thought to be highly orchestrated, they are also very complex and only a few specific mechanisms or pathways are well understood.

The visualization of our gene hierarchies typically reveals a pattern of a few coordinated modules. To explain them, we consult the Gene Ontology (GO) annotation database published by the European Bioinformatics Institute, which tags genes for participation in specific biological processes and functions or localization to specific cellular components. The GO terms belong to a separate, additional structured hierarchy tree going from general to specific. Without any further processing, the typically several thousand annotations tagging genes in the hierarchy are too numerous to be useful. So we consider only those that tag gene sets which are well-clustered along our hierarchy according to the average pairwise graph distance between nodes in the hierarchy regarded as a tree. This technique may be called tree-assisted Gene Set Enrichment Analysis. Out of the top 100 GO terms in each of the 3 basic GO categories, Process, Function, and Component, we search for the ones that might explain the apparent gene modules.

Figure 4 shows the results on the PANTHER curated gene network. Figure 5 shows the results on the GTEx lung and breast tissue RNA expression datasets. The GTEx analyses are our first examples taking advantage of the full generality of the method by involving non-synthesized node data measurements. In these examples the network topology was inferred from the sample data with a Pearson correlation cutoff, but if a prior network of interest is available it can be used instead. We caution that our experience with the usual curated networks (NetPath, PANTHER, HPRD) suggests that the choice of prior network strongly influences the results, the greater the sparsity the stronger the influence. So such results should always be compared with a control analysis based on synthetic node weights as in Figure 4.

**Figure 4:**
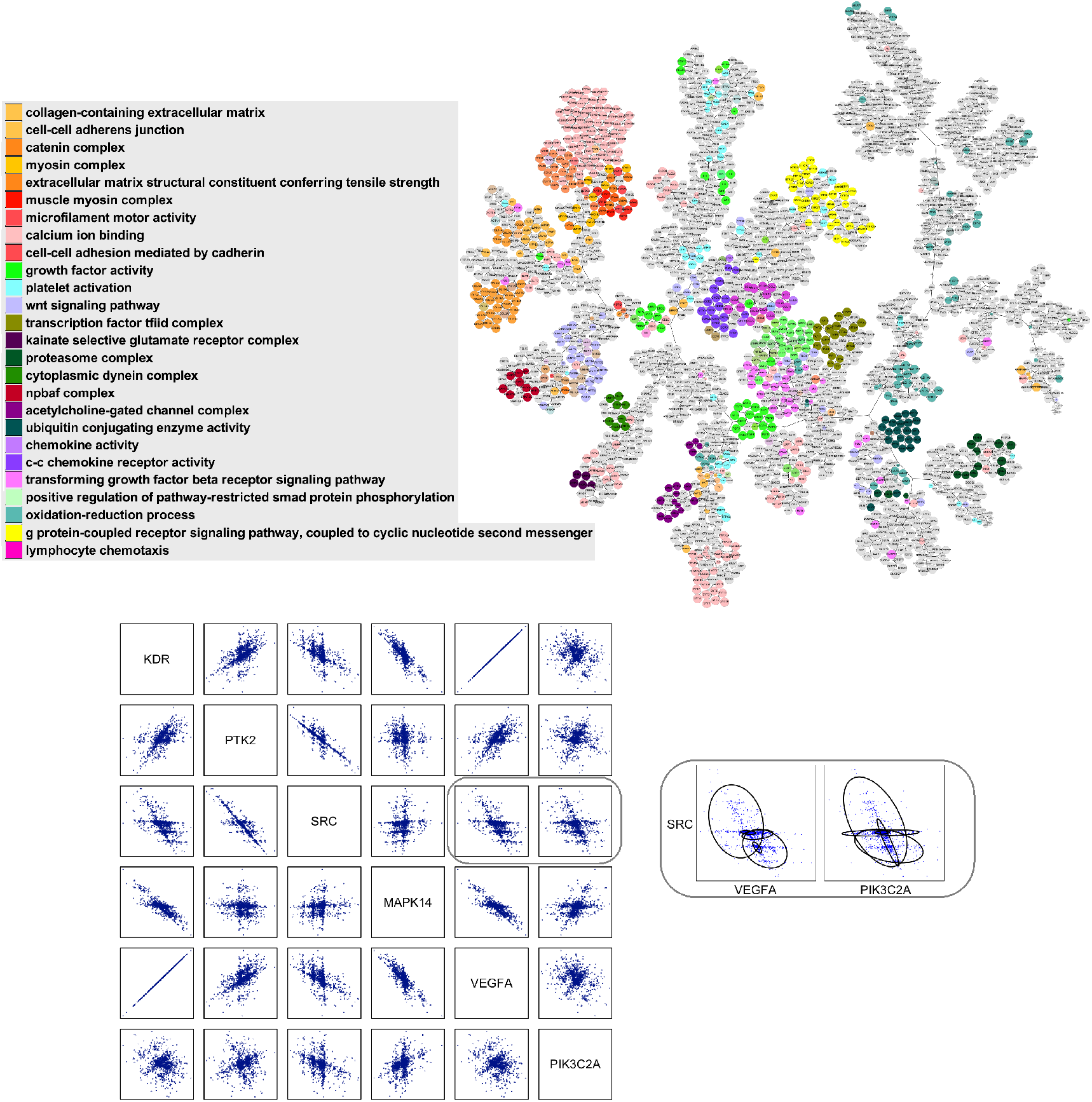
(Above) The GMT hierarchy computed from the PANTHER curated gene network with 2404 nodes and 32113 edges. Approximately 40% of the network is accounted for by well-defined processes. The gene modules appearing here are showing evidence of coordination already at the level of the network topology, without any influence from empirical node weights. This plot and GO term list could be used as a control for a separate data-driven analysis, with new annotations considered significant only when sufficiently distinct from the annotations on this list. (Below) Scatter plots of the synthesized data for selected nodes/genes, for illustration. The column containing a given gene name represents the ‘functional role profile’ of the gene with respect to the other genes. The functional role profiles of highly correlated genes are very similar (e.g. the KDR and VEGFA columns). However, similar functional roles can be observed even for uncorrelated genes. This phenomenon is one of the key differences between the GMT hierarchy and a standard correlation-based clustering. For example, the behaviors of VEGFA and PIK3C2A with respect to SRC are similar, even though the scatter plot VEGFA-PIK3C2A shows a high degree of independence. Namely, both stand in a relation of ‘inhibition’ to SRC. At a low level of our hierarchy, VEGFA and PIK3CA and perhaps other SRC inhibitors remain separate. At a middle level of our hierarchy, VEGFA and PIK3CA are closer together. In general mid level clusters may represent clusters of function, while lower levels stratify each function by the specific manner in which the function is carried out.

**Figure 5:**
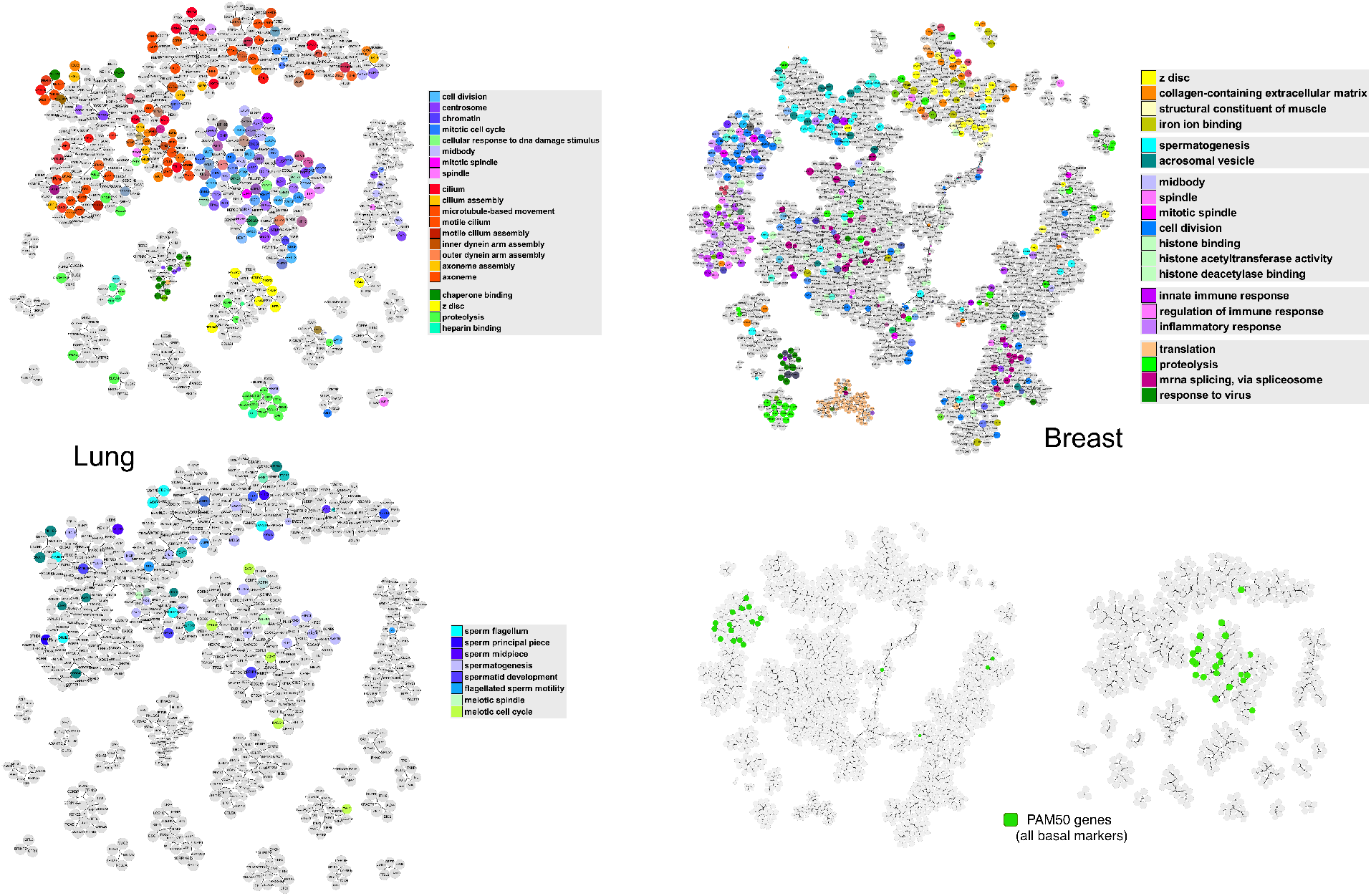
The GMT hierarchy and tree-assisted Gene Set Enrichment Analysis computed from the RNA expression profiles of the GTEx lung tissue samples (left above and below) and breast tissue samples (above right). (Below right) The 18-20 genes of the 50-gene PAM50 breast-tumor prognostic signature which appear in the breast and lung analysis turn out to be exactly the basal-marker subset identified in our earlier work [7]. Thus we find that the PAM50 is representative of only one of the key gene modules for breast tissue. Also, a substantial part of the variance in breast tumors captured by the PAM50 is likely due to variance observable already in normal (non-cancer) tissue, and moreover this type of variation is observed in normal tissues other than breast. The analysis provides evidence that the basal-marker gene submodule of the PAM50 is related to motisis, chromatin, and the architecture of the mitotic spindle (shown in shades of blue in the two upper plots). It also suggests augmentation of these markers by other genes in the apparent module, including MCM10, HMMR, ASPM, TOP2A, POLQ, RAD54L, and AURKA. Some of these genes are already known to be related to breast cancer. For example AURKA was already suggested as part of the simplified 3-gene prognostic signature for breast tumors SCMGENE [6].

## 5 Conclusion

We presented a network reduction method based on the modified optimal mass transport methodology developed for Gaussian mixture models [4]. By representing a network at several levels of complexity with respect to functional relationships between edge pairs, it was shown to be an effective method for identifying structural features in real-world datasets and benchmark networks.

## A Background on OMT for Gaussian mixture models

A Gaussian mixture model is an important instance of the general mixture model structure, a structure that is commonly utilized to study properties of populations with several subgroups [8]. Formally, a *Gaussian mixture model* (GMM) is a probability density consisting of weighted linear combination of several Gaussian components, namely

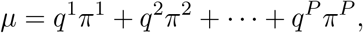

where each π^*k*^ is a Gaussian distribution and *q* = (*q*^1^, *q*^2^, …, *q^P^*)^T^ is a probability vector. Here the finite number *P* stands for the number of components of *μ*.

Let *μ*_0_, *μ*_1_ be two Gaussian mixture models of the form

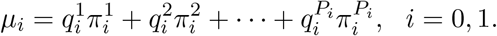

The distribution *μ_i_* is equivalent to a discrete measure *q_i_* with supports 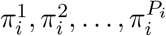 for each *i* = 0,1. The framework from [4] is based on the discrete OMT problem

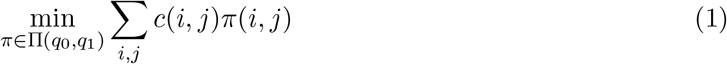

for these two discrete measures, where ∏(*q*_0_, *q*_1_) denotes the space of joint distributions with marginal distributions *q*_0_ and *q*_1_. The cost *c*(*i*, *j*) is taken to be the 2-Wasserstein metric:

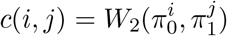

There is a closed formula for this metric:

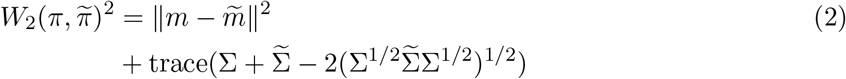

where π and 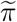 are Gaussian distributions.

The discrete OMT problem (1) always has at least one solution, and letting π* be a minimizer, we define

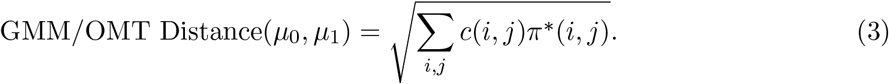

This formula, from [4], is the key formula underlying our algorithm.

**Table 1:** Notation for Algorithms 1, 2

|  |  |
| --- | --- |
| $N$ | number of nodes |
| $E$ | number of edges |
| $S$ | number of node weightings |
| $s$ | source function $[1, E] \rightarrow [1, N]$ |
| $t$ | target function $[1, E] \rightarrow [1, N]$ |
| $w$ | node weight matrix $[1, N] \times [1, S] \rightarrow \mathbb{R}$ |
| $P$ | Gaussian mixture modeling population number |
| GMM | Gaussian mixture model, ‘mclust’ R package [9] |
| $j \sim i$ | nodes $i$ and $j$ are neighbors |

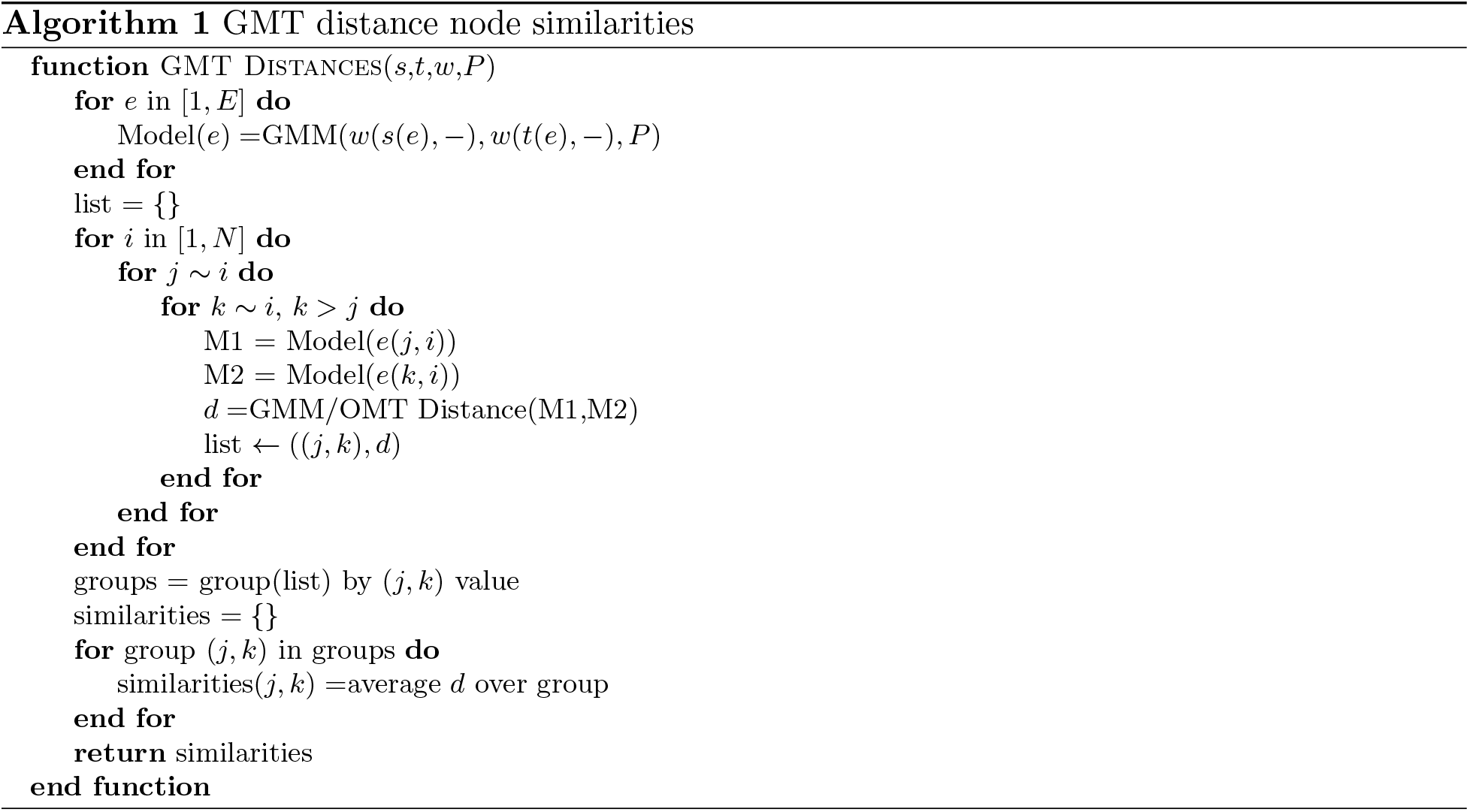

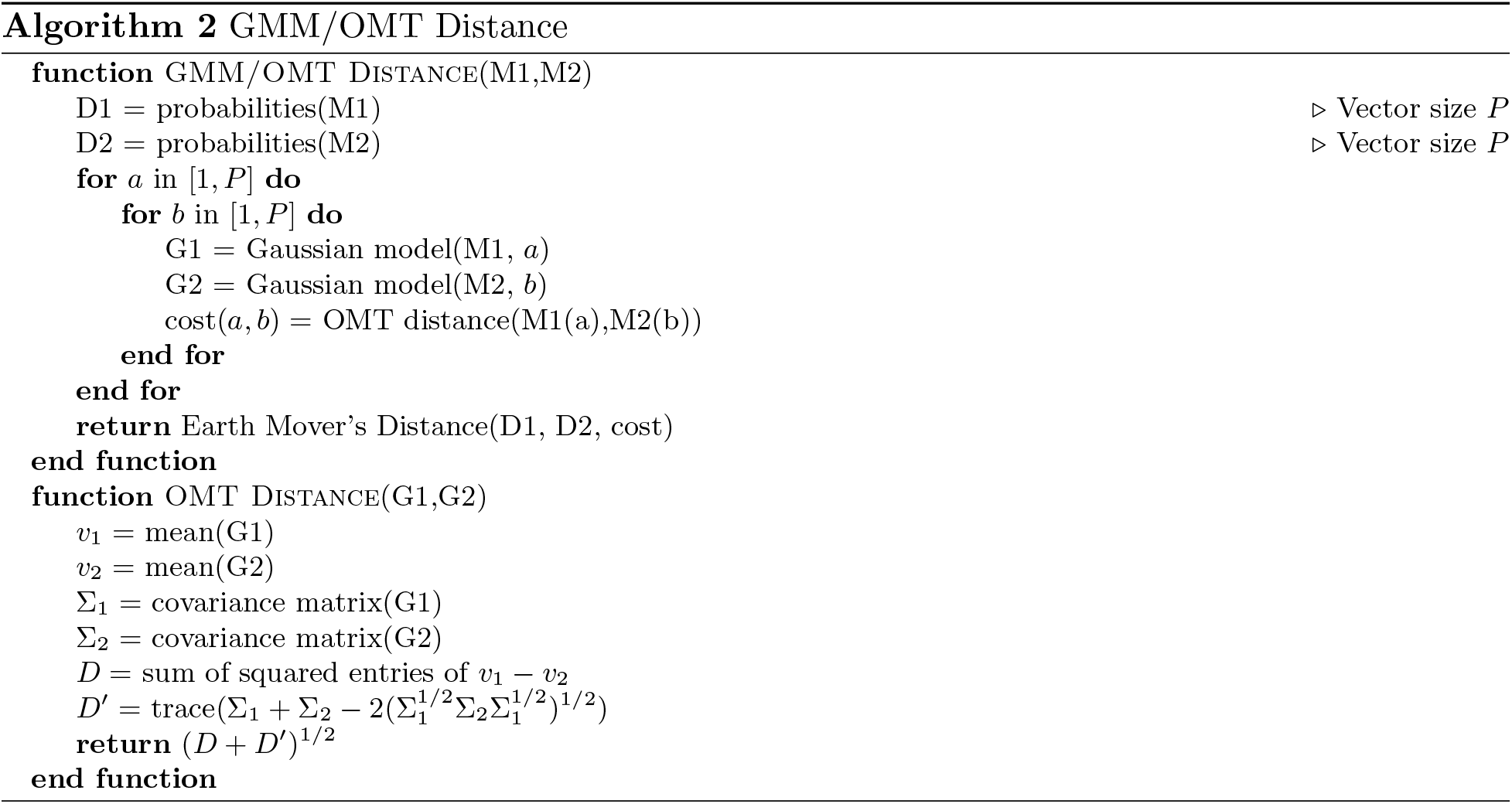

